# WEVOTE: Weighted Voting Taxonomic Identification Method of Microbial Sequences

**DOI:** 10.1101/054205

**Authors:** Ahmed A. Metwally, Yang Dai, Patricia W. Finn, David L. Perkins

## Abstract

Metagenome shotgun sequencing presents opportunities to identify organisms that may prevent or promote disease. The analysis of sample diversity is achieved by taxonomic identification of metagenomic reads followed by generating an abundance profile. Numerous tools have been developed based on different design principles. Tools achieving high precision can lack sensitivity in some applications. Conversely, tools with high sensitivity can suffer from low precision and require long computation time. In this paper, we present WEVOTE (<u>WEi</u>ghted <u>VO</u>ting <u>T</u>axonomic id<u>E</u>ntification), a method that classifies metagenome shotgun sequencing DNA reads based on an ensemble of existing methods using *k*-mer-based, marker-based, and naive-similarity based approaches. Our evaluation on fourteen benchmarking datasets shows that WEVOTE improves the classification precision by reducing false positive annotations while preserving a high level of sensitivity. WEVOTE is an efficient and automated tool that combines multiple individual taxonomic identification methods to produce more precise and sensitive microbial profiles. WEVOTE is developed primarily to identify reads generated by MetaGenome Shotgun sequencing. It is expandable and has the potential to incorporate additional tools to produce a more accurate taxonomic profile. WEVOTE was implemented using C++ and shell scripting and is available at www.bitbucket.org/ametwally/wevote

## Introduction

The microbiome plays a vital role in a broad range of host-related processes and has a significant effect on host health. Over the past decade, the culture-independent MetaGenome Shotgun (MGS) sequencing has become an emerging tool for studying the diversity and the ecology of microbial communities. One of the key steps in data analysis is the taxonomic classification of sequence reads in a metagenomic dataset.

The existing taxonomic identification methods of MGS data can be primarily classified into four categories: methods based on naive-similarity, methods based on analyzing sequence alignment results, methods based on sequence composition, such as *k*-mers, and marker-based methods. The naive-similarity-based methods rely on mapping each read to a reference database, such as the NCBI nucleotide database, and the taxonomic annotation of the best hit is assigned to the read if it passes a preset threshold. Bowtie [1], BLASTN [2], and its faster version MegaBlast [3] are the most commonly used algorithms in this category. Since the number of sequences in the database is enormous, these methods have a high probability of finding a match. Therefore, these types of methods usually achieve a higher level of sensitivity compared to other methods [4,5]. However, the major drawbacks are the increased rate of false positive annotations and the long computational time. Although it has been shown that the taxonomic profile obtained from the naive-similarity-based methods produces a large number of false positives [5,6], a vast array of researchers are still dependent on them because they do not want to sacrifice the high level of sensitivity to obtain fewer false positives annotations.

The category analyzing the results from sequence alignment includes MEGAN [7], and PhymmBL [4]. These methods consist of a preprocessing step and a post-analysis step. In MEGAN, an algorithm involving the Lowest Common Ancestor (LCA) assigns each read an NCBI taxonomic identification number (si. taxon / pl. taxa) that reflects the level of conservation within the sequence. On the other hand, PhymmBL constructs a large number of Interpolated Markov Models (IMMs) using a BLASTN query against a reference database. It subsequently computes the scores which correspond to the probability of the generated IMMs matching a given sequence. Then it classifies a read using the clade label belonging to the organism whose IMM generated the best score. The methods in this category usually require additional computational time than those in the naive-similarity methods.

The marker-based methods utilize a curated collection of marker genes where each marker gene set is used to identify a unique group of clades. The fundamental difference between these methods and the naive-similarity methods is in the reference databases. Based on how the database of the marker genes is formed, this type of methods is classified into two main subcategories: (i) methods that depend on a universal single copy marker genes database such as TIPP [8], MetaPhyler [9], and mOTU [10], and (ii) methods that depend on a clade specific marker genes database such as MetaPhlAn [11,12]. These marker-based methods can achieve high accuracy if the reads come from genomes represented by the marker gene database. Otherwise, they only achieve a low-level of sensitivity. The running time varies depending on the statistical algorithm used in each method.

The *k*-mer-based methods use DNA composition as a characteristic to achieve taxonomic annotation. The key idea is to map the *k*-mers of each read to a database of *k*-mers, and then, each read is assigned a taxonomic annotation [5,13–16]. For example, Kraken [5] uses an exact match to align the overlapped *k*-mers of the queries with a *k*-mer reference database, instead of an inexact match of the complete sequence used in the naive-similarity based methods. Because of the exact matching on short *k*-mers, many efficient data structures can be implemented for searching the *k*-mer database; thus the *k*-mer-based methods can be extremely fast. Compared to the naive-similarity methods, it was recently shown that at the genus level, *k*-mer-based methods could achieve a similar sensitivity but with higher precision [17]. However, these methods are not robust to sequences that have a high sequencing error rate because they are based on exact matching to the reference database. This limitation is demonstrated in [5]. It shows that Kraken has the lowest sensitivity compared to other methods when tested on the simBA-5 dataset.

In addition to our benchmarking,it has also revealed that different methods could generate variation in taxonomic output profiles for the same input dataset [17]. Sample type, sequencing error, and read length are the main factors that cause variation. This inconsistency in the predicted taxonomic annotations presents a challenge to investigators in the selection of identification methods and the interpretation of annotations. In this work we present a novel framework, WEVOTE (WEighted VOting Taxonomic idEntification), which takes advantage of three categories of the taxonomic identification methods; naive-similarity methods, *k*-mer-based methods, and marker-based methods. WEVOTE combines the high sensitivity of the naive similarity methods, the high precision of the *k*-mer-based methods, and the robustness of the marker-based methods to identify novel members of a marker family from novel genomes [8].

## Materials and Methods

### The WEVOTE framework and core algorithm

The core of WEVOTE is a weighting scheme organized as a taxonomic tree tallying the annotations from *N* different taxonomic identification methods. As shown in Fig. 1, the input to WEVOTE is the raw MGS reads of a microbiome sample. First, each of the *N* identification methods independently assigns a taxon for each read. If any method fails to classify the read based on the given threshold, the WEVOTE preprocessing phase assigns 0 as a taxon, indicating that the read is unclassified by the corresponding method. Then, WEVOTE identifies the taxonomic relationship of the *N* taxa per read based on the pre-configured taxonomy tree structure and casts a vote to the final taxon, which may be a common ancestor of the *N* taxa. Although the current version of our method only includes five methods, the voting scheme in our framework is flexible and allows for the inclusion or removal of different methods.

**Fig 1.**
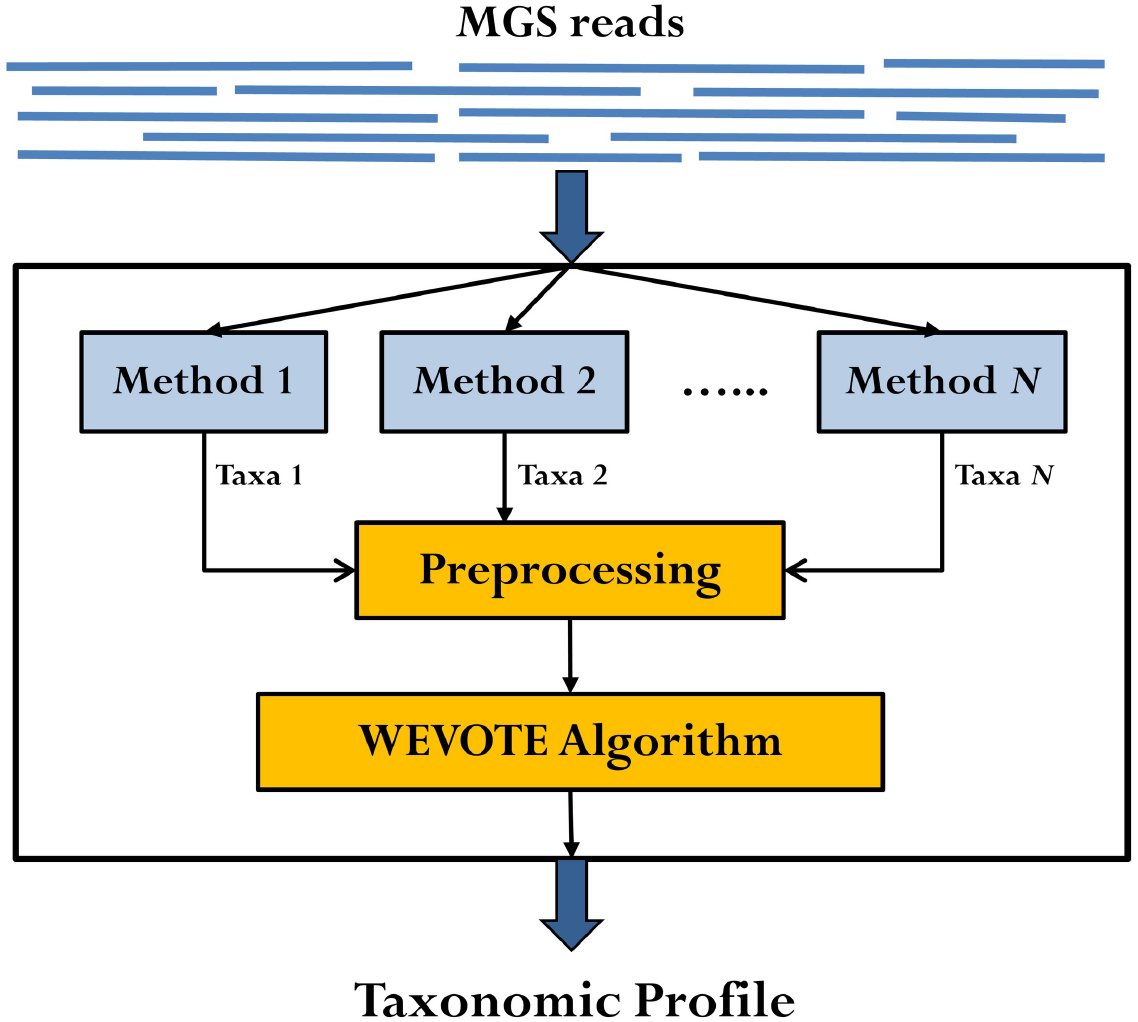
Schematic diagram of the WEVOTE framework. The input to the WEVOTE is the raw reads of the sample. First, each of the identification methods independently assigns a taxon to each read. Then, WEVOTE identifies the taxonomic relationship of the *N* taxa based on the pre-configured taxonomy tree structure and determines the final taxon assigned to each read.

WEVOTE utilizes a simplified version of the NCBI taxonomy tree as a backbone for its decision algorithm. This resolved phylogeny tree only contains the nodes that have a taxon corresponding to one of the standard taxonomic levels (Super-kingdom, Phylum, Class, Order, Family, Genus, and Species). This backbone structure facilitates and accelerates the choice of a consensus taxon based on the taxonomic annotations received from each identification tool. The decision scheme in WEVOTE is shown in Algorithm 1. Here, *N* denotes the number of tools used in the WEVOTE pipeline; *C* the number of tools that can classify the read at any taxonomic level, i.e., taxon ≠ 0; and A the number of tools that support the WEVOTE decision. The relationship *N* ⩾ *C* ⩾ *A* always holds.

#### Algorithm 1 The WEVOTE Decision Scheme

~~~
1: **procedure** WEVOTE (*N* taxa for each read)
2:    **for each** *(Read ∈ sequence file)* **do**
3:      **if** (*C* == 0) **then**
4:        *Read.Taxon* = 0
5:        *Read.DecisionScore* = 1
6:        *Read.NumSupportedTools* = N
7:     **else if** (*C* ⩾ 1) **then**
8:        *build a WeightedTree of the reported taxa*
9:        *Threshold* = *floor (C/2)*
10:        *MaxWeight* = 0
11:        *MaxNode* = 0
12:        **for each** *(Node ∈ WeightedTree and weight(Node) > Threshold)* **do**
13:          **if** *(rootToTaxon(Node) > MaxWeight)* **then**
14:            MaxWeight=rootToTaxon(Node)
15:            MaxTaxon=Node
16:          **else if** *(rootToTaxon(Node)* == *MaxWeight)* **then**
17:            MaxTaxon=LCA(Node, MaxTaxon)
18:        *Read.Taxon* = *MaxTaxon*
19:        *Read.NumSupportedTools* = *weight(Read.Taxon)*
20:        **if** (A == *C*) **then**
21:          *Read.DecisionScore* = *A/N*
22:        **else**
23:          *Read.DecisionScore* = *(A/N)* – (1/(*m * N*))
~~~

In the case that no single tool can classify the read, WEVOTE will accordingly fail to classify the read and give it a taxon 0 and score of 1. Otherwise, WEVOTE starts by building a weighted tree for each read from the taxa reported by individual tools. The weighted tree is a tree that comprises the nodes of the identified taxa along with their ancestors’ taxa including the root. The weight of any node on the weighted tree represents the number of tools that support the identification of this particular node. Next, WEVOTE annotates the read with the taxon of the node that has the highest weight from the root to that node (RootToTaxon), with the additional condition that the node itself has more weight than the WEVOTE threshold. This threshold can be set as half of the number of tools that classify a read (*C*). In the case where more than one node satisfies the WEVOTE condition, then the LCA of these nodes will be assigned as the WEVOTE decision. For each classified read, a score is also assigned to reflect the confidence of WEVOTE decision. The scoring scheme works as follows. If the number (*C*) of tools that classified the read equals the number (*A*) of tools that agreed on the WEVOTE decision, then the WEVOTE score will be calculated based on Eq. (1)

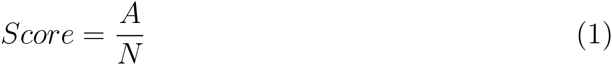

Otherwise, where *A* < *C*, the score will be calculated using Eq. (2).

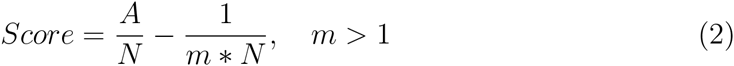

The choice of the constant m depends on how strongly one elects to penalize the disagreement among individual tools that classify the read but do not agree with the WEVOTE decision. A small value of m leads to a small WEVOTE score, implying more penalty is placed on the WEVOTE decision score, and vice versa. This scoring scheme makes the score satisfy the condition of 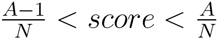. Although the score does not affect the WEVOTE decision, it would be useful if the user is interested in assessing the confidence of the taxon assignment made by WEVOTE. The default value of *m* is 2. We have chosen this value because it gives a score exactly in the middle of 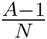 and 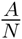. As m increases, the score skews towards the 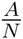 side. In order to demonstrate the decision and scoring schemes described in the WEVOTE algorithm, the case scenarios of WEVOTE for *N* = 3 are shown in Fig. 2.

**Fig 2.**
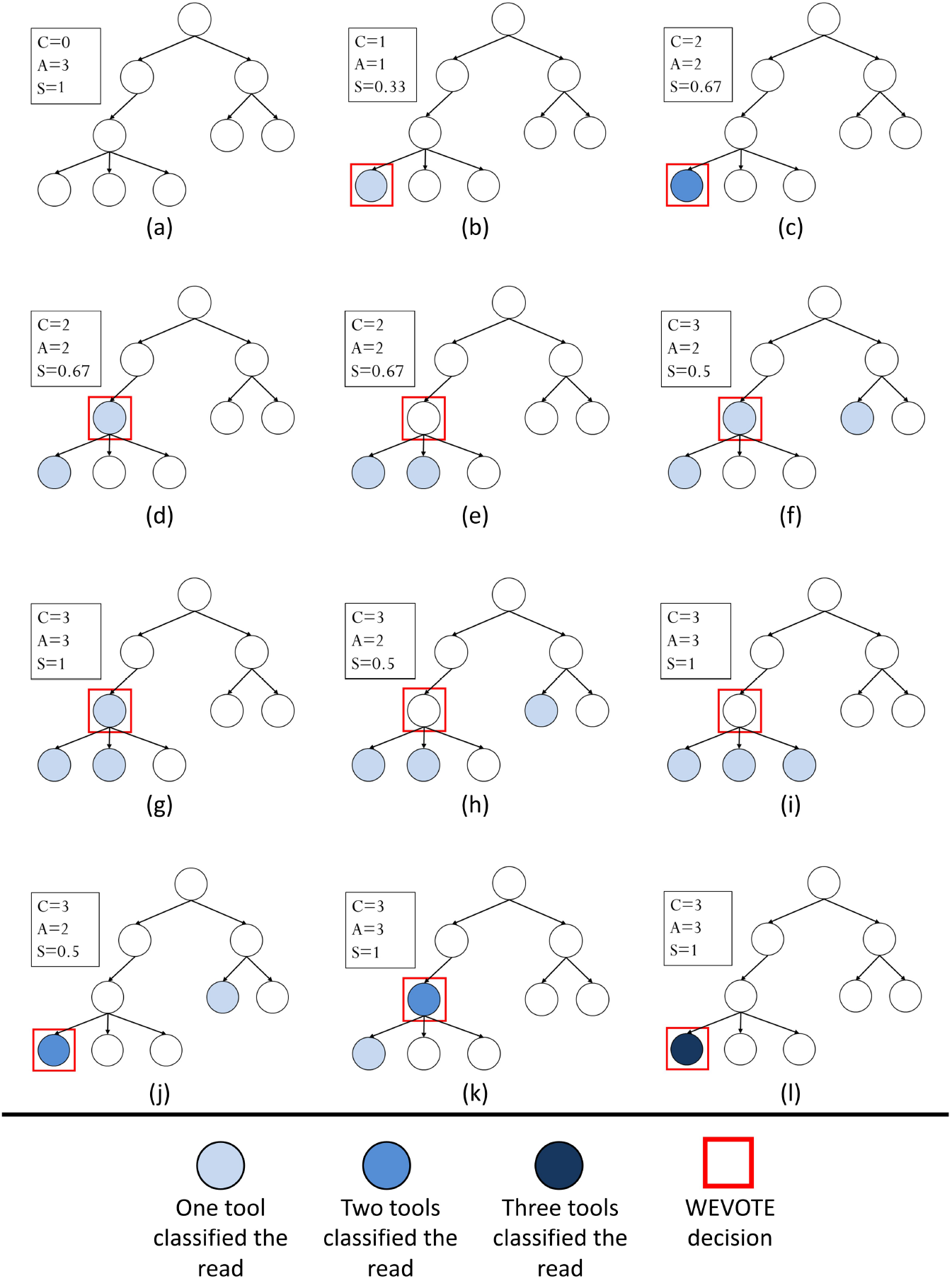
WEVOTE case scenarios using three tools. *C* denotes the # tools able to classify the read, *A* represents the # of tools that support WEVOTE Decision, and *S* represents the WEVOTE score. Scenarios are shown for (a) None of the three tools classified the read; (b) Only one tool classified the read; (c) Two tools classified the read with the same taxon; (d, e) Two tools classified the read with two different taxa; (f-i) Three tools classified the read with three different taxa; (j, k) Three tools classified the read, two taxa are identical, and the other is different; (l) Three tools identified the read with the same taxon.

### The tools used in the current implementation

In our current implementation of WEVOTE, we used BLASTN [2] to represent the naive-similarity-based methods, Kraken [5] and CLARK [13] as the identification tools representing the *k*-mer methods, and TIPP [8] and MetaPhlAn [11] representing the marker-based methods. The five tools were chosen since they are widely used and represent the three major categories of taxonomic identification methods. We selected BLASTN over MegaBlast because of its greater sensitivity. The primary reason for the increased sensitivity in BLASTN is the use of a shorter word size as a search seed. Thus, BLASTN is better than MegaBlast in finding alignments for sequences that have a sequencing error which occurs after a short length of matched bases (i.e., the initial exact match is shorter).

Kraken assigns taxonomic annotations to the reads by splitting each sequence into overlapping *k*-mers [5]. Each *k*-mer is mapped to a pre-computed database where each node in the database is the LCA taxon of all genomes that contain that *k*-mer. For each read, a classification tree is computed by obtaining all the taxa associated with the *k*-mers in that read. The number of *k*-mers mapped to each node in the classification tree is assigned as a weight for this node. The node that has the highest sum of weights from the root is used to classify the read. Kraken is an ultra-fast and highly precise algorithm for reads involving a low rate of sequencing error. CLARK is a recently released tool that is very similar to Kraken and also based on *k*-mers. It is reported to be faster and more accurate than Kraken at the genus/species level [13]. The fundamental difference between Kraken and CLARK is their backbone *k*-mers database. Kraken has only one database that can serve for the classification of metagenomic reads at any taxonomic level. If more than one genome shares the same *k*-mer, Kraken assigns this *k*-mer to their LCA taxon. CLARK, on the other hand, builds an index for each taxonomic level at which the user wishes to classify. Each level’s index has only the discriminative *k*-mers that distinguish its taxa from others.

TIPP (Taxonomic Identification and Phylogenetic Profiling) is considered a state-of-the-art tool based on a set of marker genes. It uses a customized database of 30 marker genes [18] which are mostly universal and single-copy genes. First, it performs multiple sequence alignment of each marker gene set, then builds a phylogeny tree for each marker gene and constructs a resolved taxonomy tree of these marker genes. Then, it uses SATe [19] to decompose the tree of each marker gene to many sub-trees. Subsequently, TIPP uses HMMER software [20] to build a Hidden Markov Model (HMM) for each of the sub-trees. For each query read, TIPP uses HMMER again to align the query to the HMMs. Then, TIPP uses the alignments to the HMM that have an alignment score and statistical support greater than a group of pre-set values, and places them on the precomputed taxonomic tree using pplacer [21] to assign taxonomy to the query. It has been shown that TIPP can precisely identify reads containing high sequencing error or novel members of a marker family from novel genomes [8]. The other tool chosen for this category in our implementation is MetaPhlAn. MetaPhlAn has a set of clade-specific marker genes. The marker set was built from the genomes available from the Integrated Microbial Genomes (IMG). For a given read, MetaPhlAn compares the read against the precomputed marker set using BLASTN searches in order to provide clade abundances for one or more sequenced metagenomes.

## Results and Discussion

Simulated datasets have been used in the evaluation of various taxonomic identification tools. In our assessment, we selected fourteen simulated datasets as shown in Table 1. Our choice was based on the ability of these datasets to provide the true assignment for each read rather than the true relative abundance at each taxonomic level. This information allows for the evaluation of WEVOTE based on various metrics in addition to the assessment of relative abundance.

The first three datasets were used in the evaluation of Kraken [5]. The HiSeq and MiSeq datasets are simulated from sequences obtained from non-simulated microbial projects but were sequenced using two different platforms, i.e., Illumina HiSeq and Illumina MiSeq^™^. The simBA5 is a simulated dataset with a higher percentage of error to mimic increased sequencing errors. Hence, it can be used to measure the ability of each tool to handle actual sequencing data. The simHC20 dataset was used to benchmark CLARK [13] and it contains 20 subsets of long Sanger reads from various known microbial genomes. The other ten datasets were used in MetaPhlAn [11] evaluations. HC1 and HC2 consist of reads from high-complexity, evenly distributed metagenomes that contain 100 genomes, and LC1-LC8 consist of reads from low-complexity, log-normally distributed metagenomes that contain 25 genomes. The reads from all ten MetaPhlAn datasets were sampled from KEGG v54 [22] with a length of 100 bp and an error model similar to real Illumina reads.

**Table 1.** The Benchmarking Datasets

| Source | Dataset | # reads | length (bp) | # genomes |
| --- | --- | --- | --- | --- |
| Kraken | HiSeq | 10,000 | 92 | 10 |
|  | MiSeq | 10,000 | 156 | 10 |
|  | simBA5 | 10,000 | 100 | 1,967 |
| CLARK | simHC20 | 10,000 | 951 | 20 |
| MetaPhlAn | HC1 | 999,998 | 88 | 100 |
|  | HC2 | 999,991 | 88 | 100 |
|  | LC1 | 249,995 | 88 | 25 |
|  | LC2 | 250,000 | 88 | 25 |
|  | LC3 | 250,000 | 88 | 25 |
|  | LC4 | 249,999 | 88 | 25 |
|  | LC5 | 249,999 | 88 | 25 |
|  | LC6 | 250,002 | 88 | 25 |
|  | LC7 | 250,000 | 88 | 25 |
|  | LC8 | 250,000 | 88 | 25 |

### The WEVOTE Benchmarking

Our benchmarking was performed with two variants of WEVOTE: (i) WEVOTE (*N* = 3) including BLASTN, TIPP and Kraken; and (ii) WEVOTE (*N* = 5) including BLASTN, TIPP, MetaPhlAn, Kraken, and CLARK. As described previously, BLASTN represents the naive-similarity method; TIPP and MetaPhlAn belong to the category of the marker-based methods; and Kraken and CLARK belong to the category of the *k*-mer-based methods. The default parameter values were set for the individual tools and the score penalty in WEVOTE was set at *m* = 2 (see Appendix A for full details about the commands used in the command-line). Regarding WEVOTE, we reported all results in which at least one tool supported the WEVOTE decision. With this approach, we can evaluate the accuracy of WEVOTE at the highest classification rate of the reads. By increasing the threshold, we can generate more precise results as shown later.

We first looked at how accurately each tool annotates individual reads at each taxonomic level using sensitivity and precision metrics, which are defined in Eq. (3) and Eq. (4), respectively. For each level *l* in a simulated dataset:

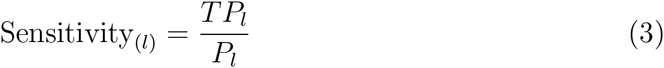

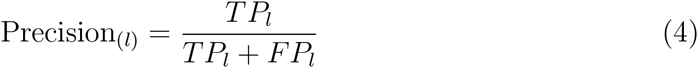

where *P_l_* denotes the number of reads annotated with some taxon at level *l* in the original dataset; *TP_l_* the number of reads correctly annotated at level *l*; and *FP_l_* the number of reads incorrectly annotated at level l.

It could be inappropriate to compare the sensitivity of all the methods used in WEVOTE, since the marker-based methods are primarily designed to calculate the microbial abundance of the sample based on the annotation of the reads that come from genes represented by the marker gene database. Based on this consideration, Fig. 3 (I) shows the sensitivity and precision of Kraken, CLARK, BLASTN, and WEVOTE; while in Fig. 3 (II), we show the precision of TIPP and MetaPhlAn separately. It is observed from Fig. 3 that WEVOTE achieves the highest level of precision and a level of sensitivity that is second only to BLASTN at the species level. At all other taxonomic levels, WEVOTE outperforms all the other individual tools in terms of sensitivity and precision in most datasets (Table S2). Note that the reason for the lower precision with *N* = 5 is because the results were reported when the minimum number of tools supported the WEVOTE decision was set at 1. If a higher level of precision is required, then the WEVOTE reporting threshold should be set at *N*/2 as explained later in Fig. 6.

**Fig 3.**
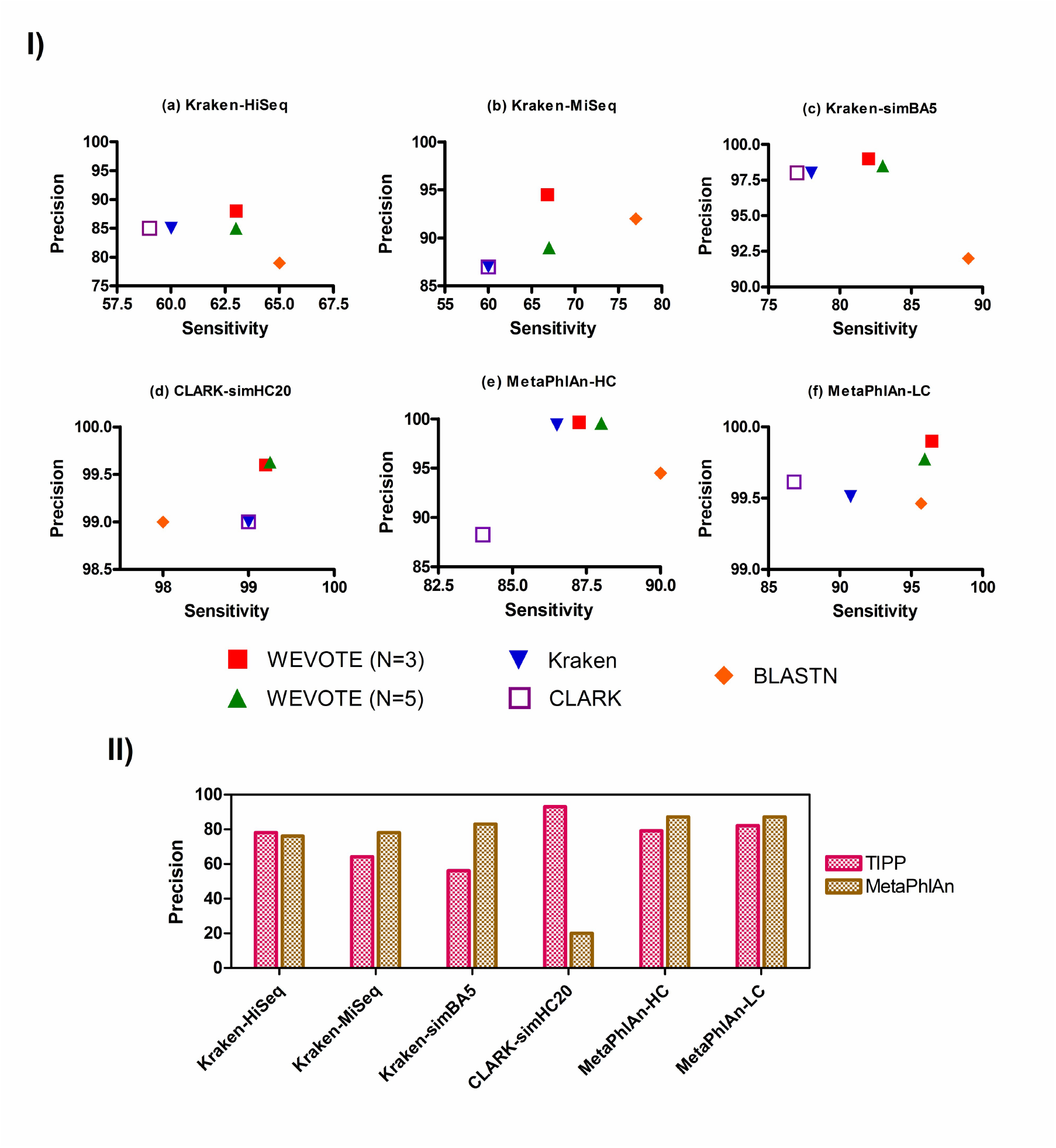
The sensitivity and precision at the species levels. sub-panel (I) shows the sensitivity and precision of tools developed to identify every read; Kraken, CLARK, BLASTN, and WEVOTE. sub-panel (II) shows the precision of marker-based tools; TIPP and MetaPhlAn. The MetaPhlAn-HC and MetaPhlAn-LC datasets are the average of two HC and eight LC datasets, respectively.

In addition, we calculated the Hellinger distance [23] (H_l_) between a sample’s metagenomic abundance profile generated by each tool and its true abundance profile at each taxonomic level *l*. The Hellinger distance measures the deviation of the predicted profile from the true profile. It is calculated as shown in Eq. (5). Here, *C_l_* is the union of all taxa that are in the true and predicted profiles at each taxonomic level *l*. For each taxon *x* at level *l, P_x_* is the predicted relative abundance and *T_x_* is the true relative abundance at taxonomic level l. The 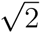 is added to the denominator to keep 0 ≤ *H_l_* ≤ 1.

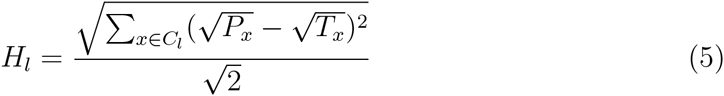

The calculation of the relative abundance (RA) differs among the tools. For tools that are developed to identify every genomic read, such as BLASTN, Kraken, and CLARK, the relative abundance is calculated as shown in Eq. (6). As mentioned before, TIPP and MetaPhlAn are not designed to identify every read. They build metagenomics abundance profile of the sample based on the annotation of the reads that come from genes represented by the marker gene database. In this case, the relative abundance of a taxon x is calculated using Eq. (7). For WEVOTE, we used Eq. (6) to calculate the RA. These two forms of relative abundance calculation are implemented in WEVOTE. It is the user option to select which method to use. However, the genomic-based method Eq. (6) is the default setting.

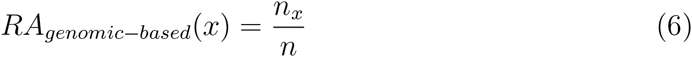

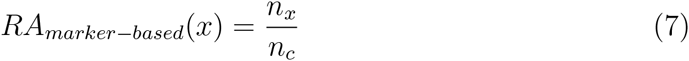

Where *n_x_* is the total number of reads classified at taxon *x, n* the total number of reads, and *n_c_* the total number of classified reads.

As the Hellinger distance represents an error distance, a small value is always preferable. Particularly, *H* = 0 means that the predicted profile is exactly the same as the true profile; while *H* = 1 means that the predicted profile is completely different from the true profile. Fig. 4 and Table S5 show the Hellinger distance between the true relative abundance profile and the profiles generated by all tools at different taxonomic levels. For all the benchmarking datasets, WEVOTE, particularly when *N* = 3, always has the smallest Hellinger distance among all other individual identification tools across all taxonomic levels. Although the Hellinger distance is marginally different for WEVOTE and BLASTN, the interpretation is quite different. The error that originates from BLASTN is due to the false positive annotations while the error that originates from WEVOTE is due to the lack of support in annotating the read at the corresponding level. TIPP and MetaPhlAn have higher Hellinger distance than other tools used in WEVOTE. This is mainly because few taxa in the datasets are predicted in low rate by them, i.e., *P_x_* being near zero for few taxa. This has led to the accumulation in the Hellinger distance. One of the reasons of the inability to predict these taxa may be because the current marker gene databases used in TIPP and MetaPhlAn do not contain sufficient markers of the genomes represented in the simulated datasets.

**Fig 4.**
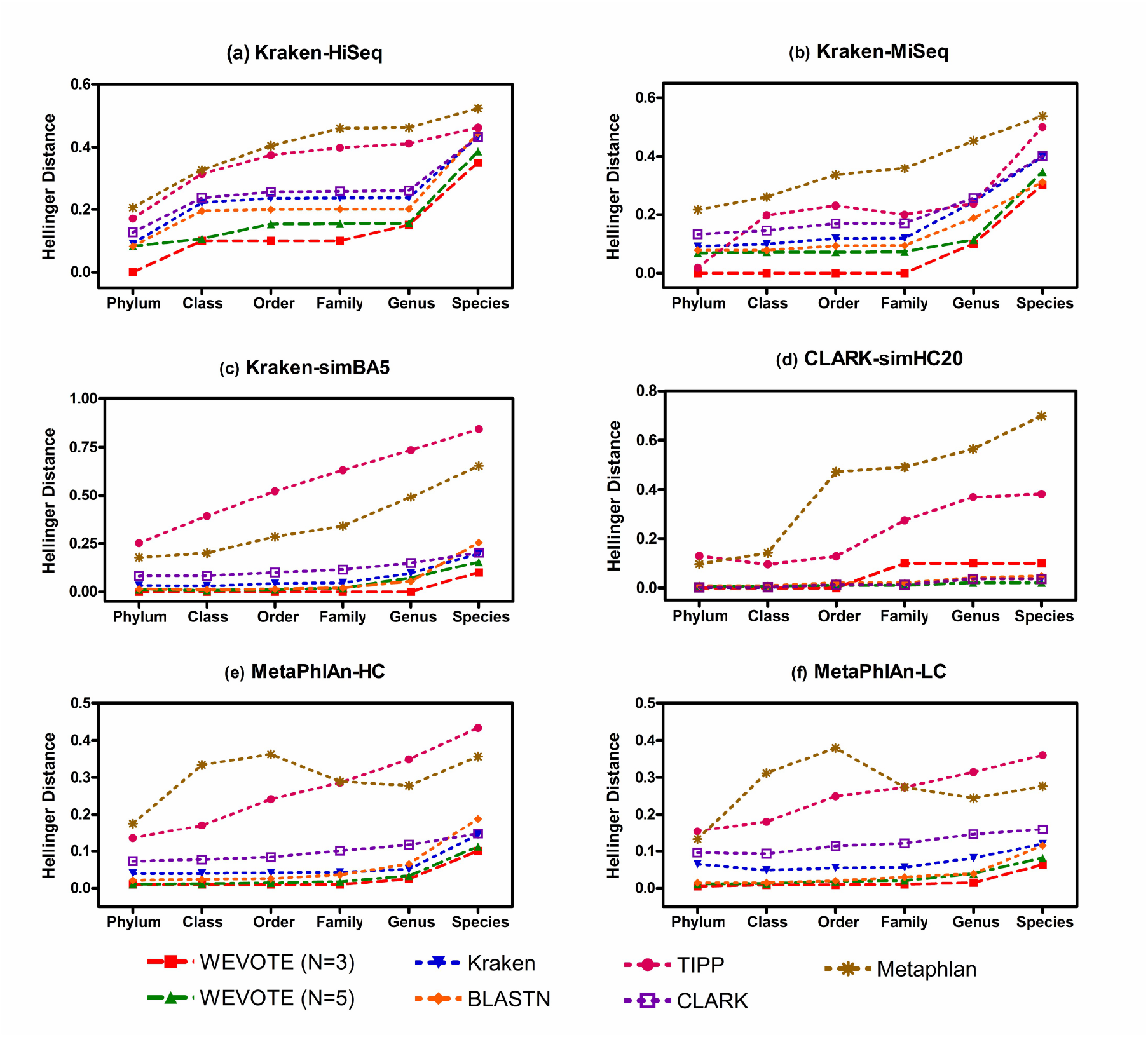
The Hellinger distance. The deviation between the predicted and the true abundance profile was measured in terms of the Hellinger distance for each tool at different taxonomic levels. Results are shown for: (a) Kraken-HiSeq dataset; (b) Kraken-MiSeq dataset; (c) Kraken-simBA5 dataset; (d) CLARK-simHC20; (e) MetaPhlAn-HC and (f) MetaPhlAn-LC. The lower the error, the more precise the corresponding tool is at the corresponding taxonomic level. *H* = 0 means that the predicted relative abundance profile is exactly the same as the true profile; while *H* =1 means that the predicted profile is completely different from the true profile.

Lastly, we examined the details of various case scenarios that were encountered in the evaluation of the two WEVOTE variants, i.e., *N* = 3 and *N* = 5. The plots in Fig. 5 show the percentages of annotations in which the individual tools support the WEVOTE decision for all the datasets. Table S4 shows the actual number of tools that support the WEVOTE decision for each dataset. It can be observed that the majority of WEVOTE annotations are determined based on more than *N*/2 agreements; 2 in the case of *N* = 3 and 3 in the case of *N* = 5. For only a small portion of each dataset, all the used tools agreed on the WEVOTE decision. An interesting observation is that a very small portion of all the classified reads by WEVOTE are in agreement with one tool when *N* = 3, or either 1 or 2 tools when *N* = 5. Therefore, if we set a threshold on WEVOTE to report the taxon at which more than half the tools are in agreement with the WEVOTE decision, then the precision of WEVOTE would increase, and its sensitivity will only be marginally decreased as demonstrated in Fig. 6. We have chosen Kraken-HiSeq and Kraken-MiSeq datasets for this investigation because they had low precision among all the used taxonomic identification tools (Fig. 3).

**Fig 5.**
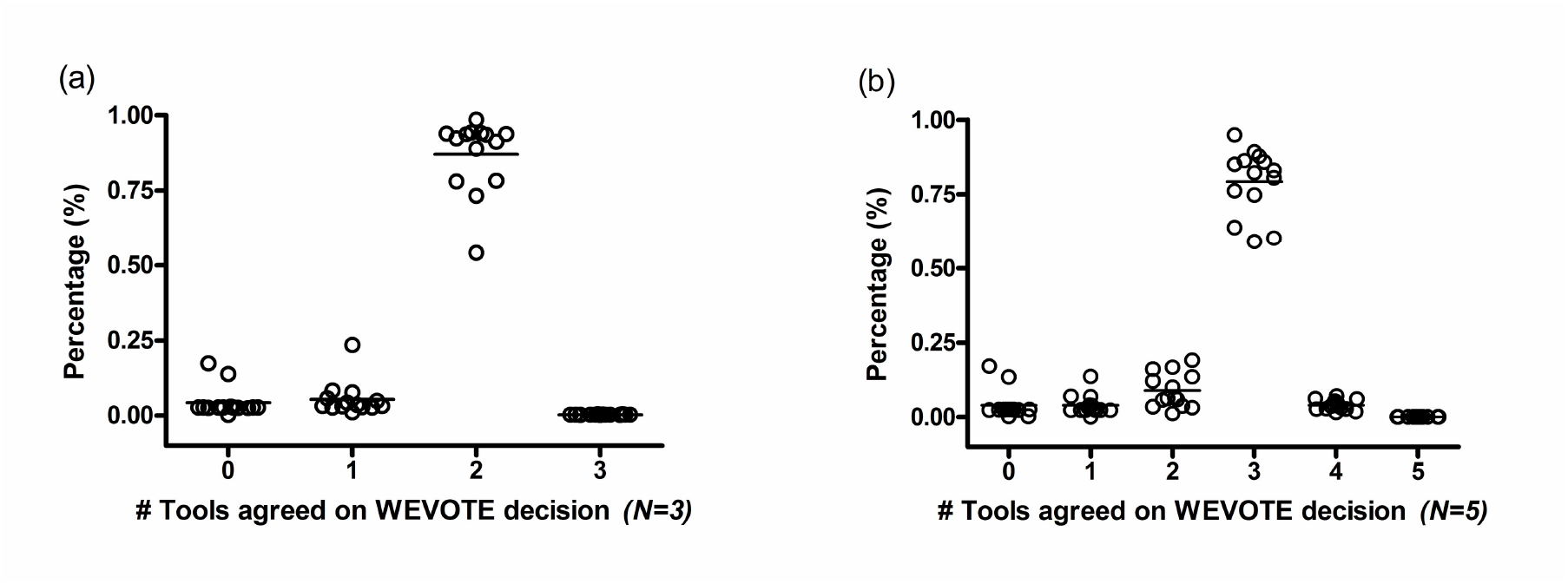
The percentage distribution of the number of individual tools that support the WEVOTE decision for the 14 datasets. Here, 0 means that the read was not classified by any tools, 1 means that one tool supports the WEVOTE assigned taxon for the read, and so on. A=3 in the case of (a) means that all the 3 tools support WEVOTE on its assigned taxon for the corresponding read, A=5 in case of (b) means that all the used 5 tools support WEVOTE on its assigned taxon for the corresponding read.

**Fig 6.**
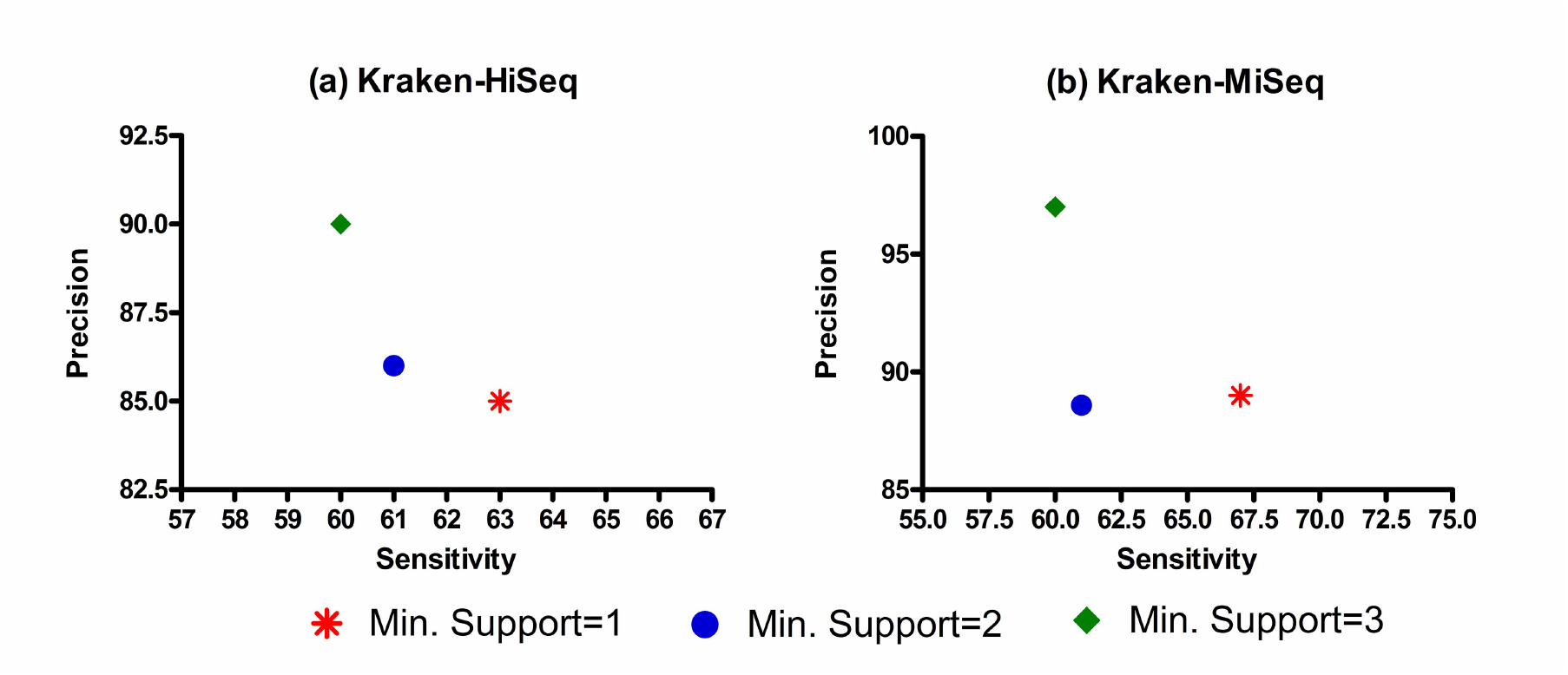
The sensitivity and precision at the species level for the WEVOTE (N=5) using different thresholds for the minimum number of tools that support the WEVOTE decision. (a) Kraken-HiSeq dataset; and (b) Kraken-MiSeq dataset.

### Computational resources and running performance

All the experiments were performed on the supercomputer (EXTREME) at the University of Illinois at Chicago. To benchmark WEVOTE, we used one node with 16 cores (Intel Xeon E5–2670 @ 2.60 GHz, cache size of 20 MB, and 128 GB RAM). Since the WEVOTE core algorithm and all the individual tools are parallelizable, we utilized 16 threads for all experiments conducted in this work. Due to the high requirement on the memory for constructing Kraken and CLARK databases, we used the Highmem node on EXTREME which has specification of 1TB RAM. In order to achieve the maximum performance from Kraken and CLARK, we used the default versions of the two tools, which require at least 80 GB of RAM. Therefore, if there is only a limited amount of memory available, users can run these tools using their mini versions, i.e., MiniKraken and CLARK-l, which only require 4 GB of RAM. In this case, the output could be 11%–25% less sensitive, but it will still preserve a high level of precision. The WEVOTE algorithm is particularly useful in this case because it can exploit the high precision level of Kraken and CLARK without using large memory machines and compensate the sensitivity by using BLASTN.

Table 2 shows the running time for each tool per dataset. For HC and LC classes of datasets, the running time is presented as the average over the datasets in each class. The standard deviation of each category is also provided (full details for all individual datasets can be found in Table S6). Kraken and CLARK finished in less than 3 minutes for any individual dataset. For BLASTN, the most time-consuming tool that is currently implemented in the WEVOTE pipeline, its running time is proportional to the number of reads and the read length in a dataset. The total time of the entire WEVOTE pipeline is the summation of the running times of the individual tools and the time needed to run the WEVOTE core algorithm. The WEVOTE core algorithm was finished execution in less than 33 seconds for any individual dataset regardless *N* = 3 or *N* = 5. The WEVOTE core algorithm is mainly affected by the number of the used tools, and more specifically, the number of tools that identified taxa for the reads. Because the running time of WEVOTE pipeline is primarily dominated by the time required by BLASTN, the pipeline running time can be reduced if many cores are used to execute BLASTN.

**Table 2.** Running time of the used tools. The time is measured in minuets.

| Simulated Dataset | Kraken | BLASTN | TIPP | CLARK | MetaPhlAn | WEVOTE Pipeline [N=5] |
| --- | --- | --- | --- | --- | --- | --- |
| HiSeq | 1 | 2 | 4 | 1 | 1 | 10 |
| MiSeq | 1 | 8 | 4 | 1 | 1 | 16 |
| simBA5 | 1 | 7 | 3 | 1 | 1 | 14 |
| simHC20 | 1 | 9 | 5 | 1 | 1 | 18 |
| HC (std) | 2 (0.0) | 30 (1.4) | 14 (0.0) | 3 (0.0) | 2 (0.0) | 53 (1.4) |
| LC (std) | 1 (0.0) | 9 (2.9) | 8 (0.5) | 2 (0.0) | 1 (0.5) | 22 (3.5) |

### Conclusion and future work

We have developed the WEVOTE framework for the consolidation of taxonomic identifications obtained from different classification tools. The performance evaluation based on the fourteen simulated microbiome datasets consistently demonstrates that WEVOTE achieves a high level of sensitivity and precision compared to the individual methods across different taxonomic levels. The major advantage of the WEVOTE pipeline is that the user can make the choice of which tools to use in order to explore the trade-off between sensitivity, precision, time, and memory. The WEVOTE architecture is flexible so that additional taxonomic tools can be easily added, or the current tools can be replaced by improved ones. Moreover, the score assigned to the taxon for each read indicates the confidence level of the assignment. This information is especially useful for the assessment of false positive annotations at a particular taxonomic level. The classification score given by WEVOTE can be used for any downstream analysis that requires the high confidence of the annotated sequences. In our current implementation, we have used a uniform weight for each method to vote. However, we will explore the potential of incorporating different weighted votes for individual methods. Future work also includes the investigation of clinical microbiome samples and experimental validation for species of interest.

## Abbreviations

WEVOTE: WEighted VOting Taxonomic idEntification method
MGS: MetaGenome Shotgun
LCA: Lowest Common Ancestor
NCBI: National Center for Biotechnology Information

## Acknowledgments

We would like to thank Dr. Nam-Phuong Nguyen and Dr. Tandy Warnow from the Department of Computer Science, The University of Illinois at Urbana-Champaign for their fruitful discussions regarding MGS taxonomic identification methods. In addition, we would like to thank Dr. Rachel Poretsky from UIC for allowing us to use the Highmem node to construct Kraken and CLARK databases. We are also grateful to Brian Nguyen and Dr. Zahraa Hajjiri for the editing of the manuscript.

## Appendix (A)

We provide information on the command line, the databases and the software version that were used for the execution of the tools: WEVOTE, BLASTN, Kraken, CLARK, TIPP, and MetaPhlAn. The complete path to each executable of database is not shown here for clarity:

### WEVOTE

The input to the WEVOTE algorithm is a CSV file. Each line of the CSV file has information about one read from the sequence fasta file. The first field of each line is the read identifier, the second field is the tool #1 taxon, the third field is tool #2 taxon, and so on. In the case of inability of any tool to classify a read, the taxon should be zero. WEVOTE, then, will use this input file along with the NCBI taxonomy database to annotate the sequences as following:

~~~
  $wevote -i <input-file.csv> -d <taxonomy-database>
-p <output-prefix> -n 16 -m 2
~~~

**Software version:** WEVOTE version 1.5.0

**Database:** NCBI taxonomy database (Downloaded on 4/17/2016).

### BLASTN

Map the reads file to the NCBI database and report the top hit:

~~~
  $blastn -db nt -query <input-file.fa> -out
<NaiveOutput> -outfmt ″6 qseqid sseqid sgi staxids length qstart qend sstart send pident evalue score
bitscore stitle″ -num_threads 16 -perc_identity 90
-max_target_seqs 1 -evalue 1e-5 -best_hit_score_edge 0.05 -best_hit_overhang 0.25
~~~

**Software version:** ncbi-blast-2.2.29+-x64-linux

**Database:** NCBI NT (Downloaded on 12/20/2015).

### Kraken

**Step 1:** Map the reads file to the Kraken database:

~~~
  $kraken --db <kraken-db> -fasta-input --threads 16
--output <KrakenOutput> [input-file.fa]
~~~

**Step 2 :** Generate Kraken report:

~~~
  $kraken-report ––db <kraken-db> <KrakenOutput> >
[KrakenOutput.report]
~~~

**Software version:** kraken-0.10.5-beta.

**Database:** used the kraken-build script to download and configure the standard Kraken database. This downloads NCBI taxonomic information, as well as the complete genomes in RefSeq for the bacterial, archaeal, and viral domains (Downloaded on 11/14/2015).

### CLARK

**Step 1:** Configure the setting and choose the database:

~~~
$set_targets.sh <CLARK-DB> bacteria viruses
~~~

**Step 2:** Map the reads file to the CLARK database:

~~~
  $classify_metagenome.sh -O <input-file.fa> -R
<output-prefix> -n 16
~~~

**Software version:** CLARKSCV1.2.3

**Database:** used the download_data.sh script to download NCBI taxonomic information, as well as the complete bacterial and virus genomes (Downloaded on 4/20/2016). In the begging of Step 2, CLARK checks whether the database exists or not, if not, it builds the default one.

### TIPP

Map the reads file to the database of 30 marker genes and reports the taxon of the classified reads:

~~~
  $run_abundance.py -f [input-file.fa] -c
/.sepp/tipp.config -x 16 -d [TIPPOutput]
~~~

**Software version:** Downloaded the source code from: https://github.com/smirarab/sepp.git on (Downloaded on 12/1/2015).

**Database:** Downloaded the references datasets from www.cs.utexas.edu/phylo/software/sepp/tipp.zip on (Downloaded on 12/1/2015).

### MetaPhlAn

Map the reads file to the database of MetaPhlAn marker genes:

~~~
  $python metaphlan.py <input-file.fa> —bowtie2db
bowtie2db/mpa ––bt2_ps sensitive-local ––bowtie2out
<output-prefix.bt2out> ––input_type multifasta ––nproc 16 > <output-file>
~~~

**Software version:** MetaPhlAn version 1.7.7

**Database:** The same marker genes database that downloaded with MetaPhlAn version 1.7.7 (Downloaded on 12/13/2015).

MetaPhlan maps each read to a clade number. The clade number is different from NCBI taxonomy ID. So, we have to use a lookup table that MetaPhlan developers provide in order to convert the clade ID to NCBI taxonomy ID.

## References

(1) Langmead B, Trapnell C, Pop M, Salzberg SL. Ultrafast and memory-efficient alignment of short DNA sequences to the human genome. Genome biology. 2009 1;10(3):R25. Available from: http://genomebiology.com/2009/10/3/R25.

(2) Altschul SF, Gish W, Miller W, Myers EW, Lipman DJ. Basic local alignment search tool. Journal of molecular biology. 1990 10;215(3):403–10. Available from: http://www.sciencedirect.com/science/article/pii/S0022283605803602.

(3) Zhang Z, Schwartz S, Wagner L, Miller W. A greedy algorithm for aligning DNA sequences. Journal of computational biology : a journal of computational molecular cell biology. 2004 1;7(1–2):203–14. Available from: http://online.liebertpub.com/doi/abs/10.1089/10665270050081478.

(4) Brady A, Salzberg SL. Phymm and PhymmBL: metagenomic phylogenetic classification with interpolated Markov models. Nature methods. 2009 9;6(9):673–6. Available from: http://dx.doi.org/10.1038/nmeth.1358.

(5) Wood DE, Salzberg SL. Kraken: ultrafast metagenomic sequence classification using exact alignments. Genome biology. 2014 1;15(3):R46. Available from: http://genomebiology.com/2014/15/3/R46.

(6) Koski LB, Golding GB. The Closest BLAST Hit Is Often Not the Nearest Neighbor. Journal of Molecular Evolution. 2001 6;52(6):540–542. Available from: http://link.springer.com/10.1007/s002390010184.

(7) Huson DH, Auch AF, Qi J, Schuster SC. MEGAN analysis of metagenomic data. Genome research. 2007 3;17(3):377–86. Available from: http://genome.cshlp.org/content/17/3/377.

(8) Nguyen NP, Mirarab S, Liu B, Pop M, Warnow T. TIPP: taxonomic identification and phylogenetic profiling. Bioinformatics (Oxford, England). 2014 12;30(24):3548–55. Available from: http://bioinformatics.oxfordjournals.org/content/30/24/3548.long.

(9) Liu B, Gibbons T, Ghodsi M, Treangen T, Pop M. Accurate and fast estimation of taxonomic profiles from metagenomic shotgun sequences. BMC genomics. 2011 1;12 Suppl 2(2):S4. Available from: http://bmcgenomics.biomedcentral.com/articles/10.1186/1471-2164-12-S2-S4.

(10) Sunagawa S, Mende DR, Zeller G, Izquierdo-Carrasco F, Berger SA, Kultima JR, et al. Metagenomic species profiling using universal phylogenetic marker genes. Nature methods. 2013 12;10(12):1196–9. Available from: http://dx.doi.org/10.1038/nmeth.2693.

(11) Segata N, Waldron L, Ballarini A, Narasimhan V, Jousson O, Huttenhower C. Metagenomic microbial community profiling using unique clade-specific marker genes. Nature methods. 2012 8;9(8):811–4. Available from: http://dx.doi.org/10.1038/nmeth.2066.

(12) Truong DT, Franzosa EA, Tickle TL, Scholz M, Weingart G, Pasolli E, et al. MetaPhlAn2 for enhanced metagenomic taxonomic profiling. Nature Methods. 2015 9;12(10):902–903. Available from: http://dx.doi.org/10.1038/nmeth.3589.

(13) Ounit R, Wanamaker S, Close TJ, Lonardi S. CLARK: fast and accurate classification of metagenomic and genomic sequences using discriminative k-mers. BMC Genomics. 2015 3;16(1):236. Available from: http://bmcgenomics.biomedcentral.com/articles/10.1186/s12864-015-1419-2.

(14) Ames SK, Hysom DA, Gardner SN, Lloyd GS, Gokhale MB, Allen JE. Scalable metagenomic taxonomy classification using a reference genome database. Bioinformatics (Oxford, England). 2013 9;29(18):2253–60. Available from: http://bioinformatics.oxfordjournals.org/content/29/18/2253.short.

(15) Menzel P, Lee Ng K, Krogh A. Kaiju: Fast and sensitive taxonomic classification formetagenomics; 2015. Available from: http://biorxiv.org/content/early/2015/12/18/031229.abstract.

(16) Rosen GL, Reichenberger ER, Rosenfeld AM. NBC: the Naive Bayes Classification tool webserver for taxonomic classification of metagenomic reads. Bioinformatics (Oxford, England). 2011 1;27(1):127–9. Available from: http://bioinformatics.oxfordjournals.org/content/27/1/127.

(17) Lindgreen S, Adair KL, Gardner PP. An evaluation of the accuracy and speed of metagenome analysis tools. Scientific Reports. 2016 1;6:19233. Available from: http://www.nature.com/srep/2016/160118/srep19233/full/srep19233.html.

(18) Wu M, Eisen JA. A simple, fast, and accurate method of phylogenomic inference. Genome Biology. 2008 1;9(10):R151. Available from: http://genomebiology.com/2008/9/10/R151.

(19) Liu K, Warnow TJ, Holder MT, Nelesen SM, Yu J, Stamatakis AP, et al. SATe-II: very fast and accurate simultaneous estimation of multiple sequence alignments and phylogenetic trees. Systematic biology. 2012 1;61(1):90–106. Available from: http://www.ncbi.nlm.nih.gov/pubmed/22139466.

(20) Eddy SR. Profile hidden Markov models. Bioinformatics (Oxford, England). 1998 1;14(9):755–63. Available from: http://www.ncbi.nlm.nih.gov/pubmed/9918945.

(21) Matsen FA, Kodner RB, Armbrust EV. pplacer: linear time maximum-likelihood and Bayesian phylogenetic placement of sequences onto a fixed reference tree. BMC bioinformatics. 2010 1;11:538. Available from: http://www.pubmedcentral.nih.gov/articlerender.fcgi?artid=3098090{&}tool=pmcentrez{&}rendertype=abstract.

(22) Kanehisa M, Goto S, Furumichi M, Tanabe M, Hirakawa M. KEGG for representation and analysis of molecular networks involving diseases and drugs. Nucleic acids research. 2010 1;38(Database issue):355–60. Available from: http://www.pubmedcentral.nih.gov/articlerender.fcgi?artid=2808910{&}tool=pmcentrez{&}rendertype=abstract.

(23) Deza E, Deza MM. Encyclopedia of Distances. Berlin, Heidelberg: Springer Berlin Heidelberg; 2009. Available from: http://link.springer.com/10.1007/978-3-642-00234-2.

